# RNAbpFlow: Base pair-augmented SE(3)-flow matching for conditional RNA 3D structure generation

**DOI:** 10.1101/2025.01.24.634669

**Authors:** Sumit Tarafder, Debswapna Bhattacharya

**Affiliations:** Department of Computer Science, Virginia Tech, Blacksburg, Virginia, 24061, USA

**Keywords:** RNA 3D structure modeling, generative modeling, deep learning, flow matching

## Abstract

Despite the groundbreaking advances in deep learning-enabled methods for biomolecular modeling, predicting accurate three-dimensional (3D) structures of RNA remains challenging due to the highly flexible nature of RNA molecules combined with the limited availability of evolutionary sequences or structural homology. Here we introduce RNAbpFlow, a novel sequence- and base-pair-conditioned SE(3)-equivariant flow matching model for generating RNA 3D structural ensemble. Leveraging a nucleobase center representation, RNAbpFlow enables end-to-end generation of all-atom RNA structures without the explicit or implicit use of evolutionary information or homologous structural templates. Experimental results show that base pairing conditioning leads to broadly generalizable performance improvements over current approaches for RNA topology sampling and predictive modeling in large-scale benchmarking. RNAbpFlow is freely available at https://github.com/Bhattacharya-Lab/RNAbpFlow.

## 1 Introduction

The determination of three-dimensional (3D) structures of RNA has become a crucial challenge in structural biology, driven by the growing interest in RNA-based therapeutics [1, 2]. High-resolution characterization of 3D RNA structure is essential for the design and understanding of RNA molecules with specific therapeutic functions [3], thus expanding the scope of RNA-mediated drug discovery [4]. However, the intrinsic conformational flexibility of RNA presents significant challenges for experimental structure determination methods such as X-ray crystallography, nuclear magnetic resonance (NMR) spectroscopy, and cryo-electron microscopy (Cryo-EM). Computational RNA structure prediction, therefore, is emerging as an attractive alternative to fill the gap in RNA structure space and to elucidate RNA conformational dynamics that underpins diverse cellular processes [5].

Traditional RNA 3D structure prediction methods include template-based approaches like ModeRNA [6] and RNAbuilder [7], which rely on homologous structural information, as well as physics- and/or knowledge-based methods such as FARFAR2 [8], 3dRNA [9], RNAComposer [10], and Vfold3D [11], which exploit biophysical potentials and pre-built fragment libraries to assemble full-length RNA structures. However, these approaches are constrained by the scarcity of RNA structural data in the PDB [12] and are often computationally prohibitive, making them less suitable for the prediction of large RNAs with complex topologies [13, 14]. Although physics-based methods combined with expert human intervention have demonstrated success in community-wide blind RNA-Puzzles [15] and CASP (Critical Assessment of Structure Prediction) challenges [16], there remains a critical need for fully automated, fast and accurate methods for computational modeling of RNA structures.

Inspired by the transformative impact of AlphaFold 2 [17] on protein structure prediction [18], a growing number of deep learning-based methods have recently been developed for modeling RNA structures including DRfold [13], trRosettaRNA [14], trRosettaRNA2 [19], RoseTTAFoldNA [20], RhoFold+ [21], and NuFold [22] by leveraging attention-powered transformer architectures [23]. However, except for DRfold, most of these methods are highly dependent on explicit evolutionary sequence information derived from multiple sequence alignments (MSA) or implicitly make use of homologous information learned by biological language models [21]. Obtaining reliable MSAs for RNA sequences poses significant challenges due to the isosteric nature of base pair interactions, hindering sequence alignment efforts [24]. Furthermore, many existing methods fail to fully leverage the RNA base pair (2D) information, including canonical and non-canonical base pairing interactions, key determinants of the final 3D conformation of RNA [25, 26]. Finally, the static structure predictions made by these approaches might be inadequate to capture the inherent conformational flexibility of an RNA molecule that often adopts a distribution of conformational states instead of folding into a static structure [5, 27]. Thus, there is an urgent need to develop improved computational methods that can generate a conformational ensemble of all-atom RNA 3D structures directly from the nucleotide sequence by making use of base pairing information without explicitly or implicitly using any evolutionary information.

Diffusion-based generative modeling has achieved remarkable success in the image domain and is now attracting significant attention in structural bioinformatics, most notably in AlphaFold 3 [28] for biomolecular interaction prediction, as well as 3D protein backbone generation with approaches such as RFdiffusion [29] and FrameDiff [30]. More recently, methods such as FrameFlow [31] have showcased the power of SE(3)-equivariant flow matching, achieving better precision in designability than their diffusion-based counterpart but with reduced sampling costs. RNA-FrameFlow [32], a recent adaptation of FrameFlow for RNA, is the first generative model that specifically targets 3D RNA backbone generation. However, the method employs naïve unconditional flow matching, generating backbones without using sequence or base pairing information. Such a limitation highlights the opportunity to develop improved deep generative models for RNA that can efficiently sample a large conformational ensemble of all-atom RNA 3D structures by explicitly conditioning on the nucleotide sequence and base pairing information through conditional flow matching, thereby offering an elegant combination of principles-based and data-driven approach to RNA structural ensemble generation free from sequence- and structural-level homology.

Here we present RNAbpFlow, a novel sequence- and base-pair-conditioned SE(3)-equivariant flow matching model for generating all-atom RNA conformational ensemble. The major contributions of this work lie in the methodology development of the first conditional flow matching-based method for RNA 3D structure generation. First, RNAbpFlow incorporates conditions on the nucleotide sequence and base pairing information from three complementary base pair annotation methods to comprehensively capture canonical and non-canonical interactions. Second, by incorporating a nucleobase center representation that enables the optimization of angles of all rotatable bonds of nucleobases, RNAbpFlow directly outputs all-atom RNA structures in an end-to-end fashion, bypassing the need for a post hoc geometry optimization module which is impractical in the context of large-scale sample generation. Third, we introduce base pair-centric auxiliary loss functions to enable maximal realization of the canonical and non-canonical base pairing interactions. We empirically observe performance improvements when base pairing condition is introduced across a wide range of evaluation metrics, outperforming a recent Molecular Dynamics (MD) simulation-based global topology sampling method RNAJP [33] which explicitly considers base pairing and base stacking interactions. Additionally, RNAbpFlow generalizes well for sequence- and base-pair-conditioned RNA 3D structure prediction compared to several state-of-the-art methods, including top-performing automated servers participating in recently concluded CASP16 challenge. RNAbpFlow is freely available at https://github.com/Bhattacharya-Lab/RNAbpFlow.

## 2 Results

### 2.1 Overview of the RNAbpFlow framework

An overview of our method, RNAbpFlow, is shown in **Figure 1**. Our framework is built on the foundations of FrameFlow [31], a flow matching formulation tailored for fast protein backbone generation on the SE(3) frame representation. To represent each nucleotide in an RNA sequence as a rigid body frame defined by a translation from the global origin and a rotation matrix, as well as for the full atomic RNA 3D structure generation in an end-to-end manner, we follow the nucleotide representation presented in NuFold [22]. The rotation matrix is constructed using the Cartesian coordinates of C1^*′*^ as the origin of the local frame and O4^*′*^– C1^*′*^ – C2^*′*^ for orientation. Given an RNA sequence of length *N*, we begin with *N* such frames sampled from a Gaussian distribution as the starting point for our iterative sampling process. Our method incorporates conditions on the nucleotide sequence and base pairing information to guide the conformational sampling process to generate the all-atom RNA structure. The three base-frame atoms: O4^*′*^, C1^*′*^, and C2^*′*^ are derived from the learned frame representation, while the first nitrogen of the base (N1 for pyrimidines, N9 for purines) is imputed using tetrahedral geometry. The remaining atoms are partitioned into ten frames, and the corresponding atomic coordinates are generated based on the iterative update of the frames using the 9 predicted torsion angles lying on the bonds that connect these frames, leading to an all-atom RNA structure.

**Figure 1.**
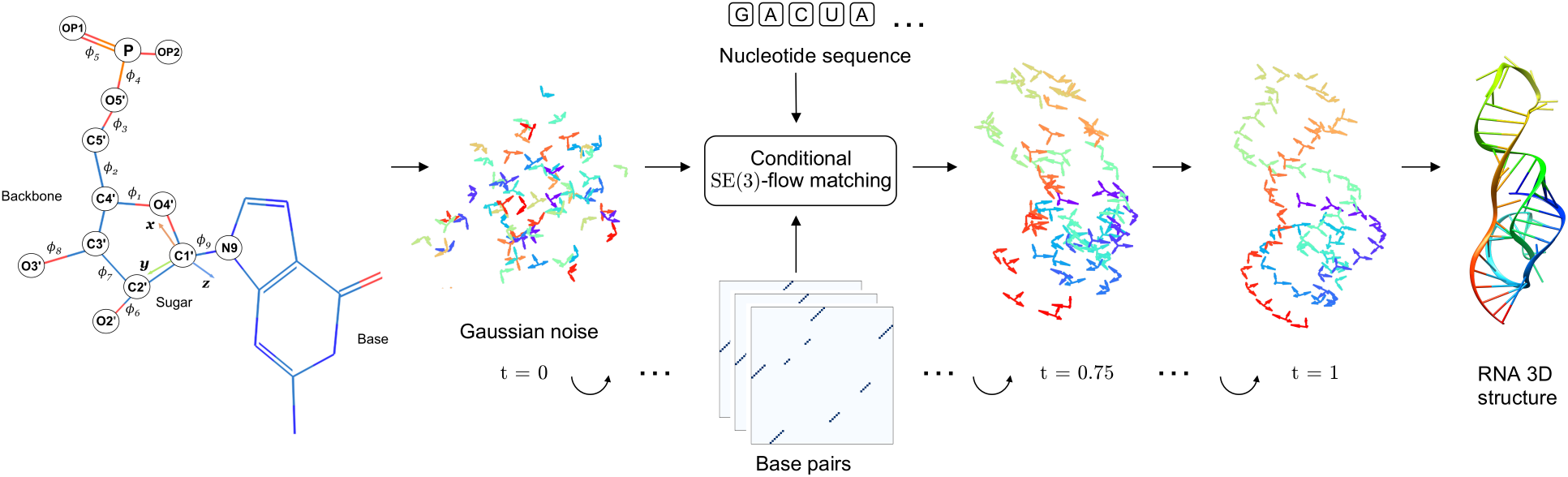
Overview of RNAbpFlow. RNAbpFlow is a sequence- and base-pair-conditioned SE(3)-equivariant flow m atching m odel f or g enerating R NA 3 D s tructural e nsemble. U sing the n ucleotide s equence a nd b ase pairing information as conditions, our end-to-end framework enables efficient sampling of all-atom RNA 3D structures based on a nucleobase center representation without explicitly or implicitly using any evolutionary information.

### 2.2 Experimental setup and performance evaluation

For training and performance evaluation, we utilize two existing and publicly available training sets, each tailored for a specific evaluation benchmark. We explicitly ensure that there is no redundancy between any training set and its corresponding test set as well as CASP15 and CASP16 targets. To develop and internally benchmark our method, we rely on the train-test split provided by RNA3DB [34], a recently released RNA dataset specifically curated to eliminate both sequence and structural redundancy—making it an ideal for training deep learning models. To ensure a fair evaluation on CASP15 targets and avoid any potential data leakage, we deliberately avoid using RNA3DB for training in this case. Instead, we curate a completely new and separate training set from trRosettaRNA [14] which only includes RNA chains released before January 2022 in the Protein Data Bank (PDB) [12]. This cutoff predates the release of the first CASP15 target in May 2022, ensuring that none of the CASP15 structures were accessible during training, maintaining complete non-redundancy following the approach of trRosettaRNA [14]. For CASP16, we curate a training set entirely from RNA3DB which is augmented with a cross-distillation set curated from bpRNA-1m (90) [35]. Details of the further curation process of these training and benchmark sets are provided in Section 4.4. To evaluate the sampling performance of RNAbpFlow, we compare against RNAJP [33], a recent coarse-grained molecular dynamics simulation-based method that samples RNA 3D structures from base pairing input. For assessing the predictive performance, we benchmark RNAbpFlow against deep learning-based methods including AlphaFold 3 [28], trRosettaRNA2 [19], DRfold [13] and RhoFold+ [21], and physics- or knowledge-based approaches such as RNAComposer [10], 3dRNA [36], and Vfold-Pipeline [37], all of which operate on sequence and base pairing input. Evaluation metrics span global fold similarity and local structural accuracy, including TM-score [38], GDT-TS [39], lDDT [40], RMSD, INF-All [41] and clash score [42]. Details of the evaluation process is discussed in Section 4.5.

### 2.3 Impact of base pairs on RNA 3D generation

To assess the significance of the base pairing information incorporated in our conditional flow matching formulation for RNA 3D structure generation, we individually condition on base pairs extracted from the native 3D structures using each of the three different 2D annotation tools described in Section 4.1 as well as their combination. We independently train each variant on the RNA3DB training set (detailed in Section 4.4) and evaluate on the 52 targets from the corresponding non-redundant test set. We also train a baseline model by conditioning only on the nucleotide sequence but not on the base pairing information. For each target sequence, we generate 1,000 3D structural samples and calculate the maximum and mean of both TM-score and lDDT. The distribution of the average of these scores across all 52 targets is shown in **Figure 2** based on various base pair conditioning (or lack thereof).

**Figure 2.**
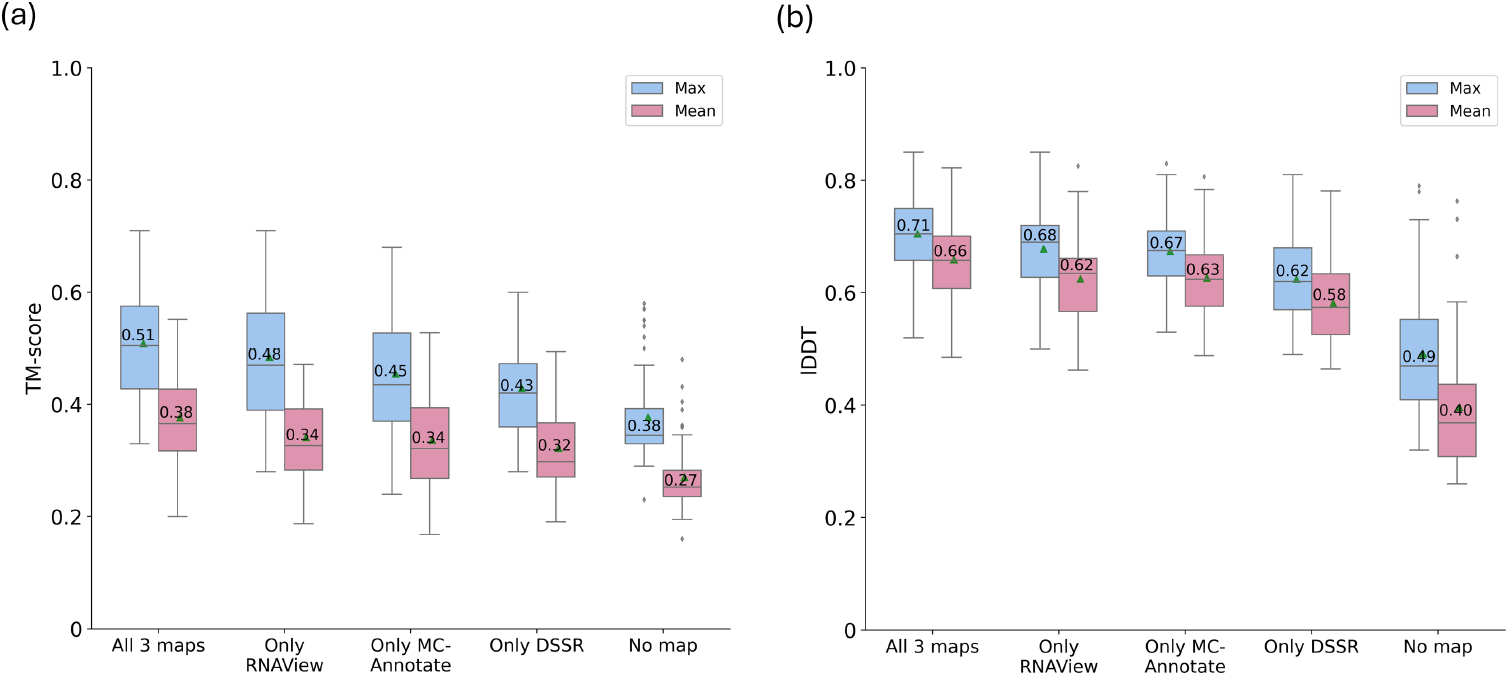
Distributions of the maximum and mean of 1000 3D structural samples per target in terms of (a) TM-score and (b) lDDT across 52 RNA3DB test targets for various base pairing conditioning (or lack thereof). The green triangles indicate averages.

As shown in **Figure 2**, RNAbpFlow achieves the best performance when all three base pair maps are incorporated as condition, resulting in an average over the maximum per-target TM-score distribution of 0.51 and lDDT of 0.71. This represents an average increase of 34.2% in TM-score and 44.9% in lDDT compared to the baseline sequence-conditioned variant of our flow matching formulation, which achieves an average over the maximum per-target TM-score distribution of 0.38 and lDDT of 0.49. These improvements underscore the critical role of incorporating base pair information in RNA 3D generative modeling. The relative contribution of individual base pair maps is also evident from **Figure 2**, which demonstrates that none of the individual maps can alone outperform their combination. Furthermore, the distributions reveal the consistency of the generated sample qualities when all three maps are used, which achieves a relatively smaller interquartile range while maintaining the highest mean values for both TM-score and lDDT distributions, indicating both high-quality samples and reduced variability. This consistency is also reflected in sampling performance where RNAbpFlow successfully generates at least one 3D structure with a TM-score exceeding 0.45 (a threshold for assessing RNA global fold correctness [43]) for 37 out of 52 targets with 71.2% correctly folded targets from the test set, demonstrating the robustness of RNAbpFlow in RNA 3D structure generation when accurate base pairing information is available.

### 2.4 Structural ensemble generation performance

To evaluate the sampling performance of RNAbpFlow, we compare against RNAJP, a recent coarse-grained MD simulation-based method for RNA 3D structure sampling with explicit consideration of base pair information, including non-canonical base pairing and base stacking interactions as well as long-range loop–loop interactions. For the benchmark set of 12 RNA targets containing three-way junctions described in Section 4.4, we generate 1,000 3D structural samples per target and compute the mean and maximum scores (TM-score and lDDT) for each target, which are then averaged across the 12 RNAs. To ensure a fair comparison, we randomly select 1,000 decoy structures for each target from the entire ensemble provided by RNAJP.

The results, summarized in **Table 1**, demonstrate that RNAbpFlow consistently outperforms RNAJP in both metrics. For example, RNAbpFlow achieves an average over the mean lDDT score of 0.67, surpassing that of 0.59 achieved by RNAJP. Similarly, in terms of global topology sampling, RNAbpFlow generates higher-quality structures, achieving an average over the mean TM-score of 0.38 compared to RNAJP’s 0.32. To evaluate sampling efficacy, we calculate the fraction of targets for which at least one correct fold (TM-score *>* 0.45 or lDDT *>* 0.75) appears in the ensemble [43] as well as the fraction of all decoys that are correctly folded. RNAbpFlow finds a TM-score–based correct fold in 50% of the RNAs and an lDDT-based correct fold in 25%, compared to RNAJP’s 41.67% and 0.0%, respectively. Across all 12, 000 decoys generated by RNAbpFlow, 17.3% achieve TM-score *>* 0.45 and 15.0% achieve lDDT *>* 0.75. By contrast, only 1.73% of RNAJP’s decoys exceed TM-score *>* 0.45, and 0.0% exceed lDDT *>* 0.75. These results demonstrate that RNAbpFlow not only outperforms RNAJP in terms of top-scoring structures but also results in a substantially higher proportion of high-quality decoys, highlighting its efficiency in sampling both global topology and local conformations.

**Table 1:** Sampling results in terms of maximum and mean scores of 1,000 3D structural samples for 12 RNAJP test RNAs. Values in bold indicate the best performance.

| Method | TM-score |  | IDDT |  |
| --- | --- | --- | --- | --- |
|  | Max | Mean | Max | Mean |
| RNAbpFlow | **0.5** | **0.38** | **0.72** | **0.67** |
| RNAJP | 0.45 | 0.32 | 0.65 | 0.59 |

**Figure 3** showcases two representative examples from the RNAJP benchmark set, illustrating the predictive performance of RNAbpFlow compared to the RNAJP method. For each target, the best structure from the 3D structural samples generated by both methods is selected based on the highest TM-score. For the first case study target RNA 2HGH of 55 nucleotides containing one three-way junction, the b est structure predicted by RNAbpFlow shown in **Figure 3(a)** achieved a TM-score of 0.58, indicating a correct global fold and a precise prediction of the three-way junction, with an all-atom RMSD of 2.58 Å. By contrast, the RNAJP method achieves a lower TM-score of 0.42, partly due to the misorientation of the three-way junction and the hairpin loop. A visual inspection of the predicted base pairs extracted from the 3D structures in **Figure 4(a)** demonstrates that RNAbpFlow more effectively preserves the overall base pairing interactions, particularly the non-canonical ones in the three-way junction region than RNAJP. By capturing the non-canonical base pair interactions with high fidelity, RNAbpFlow achieves a higher INF-NWC score of 0.78, which is much higher than that of RNAJP (0.52).

**Figure 3.**
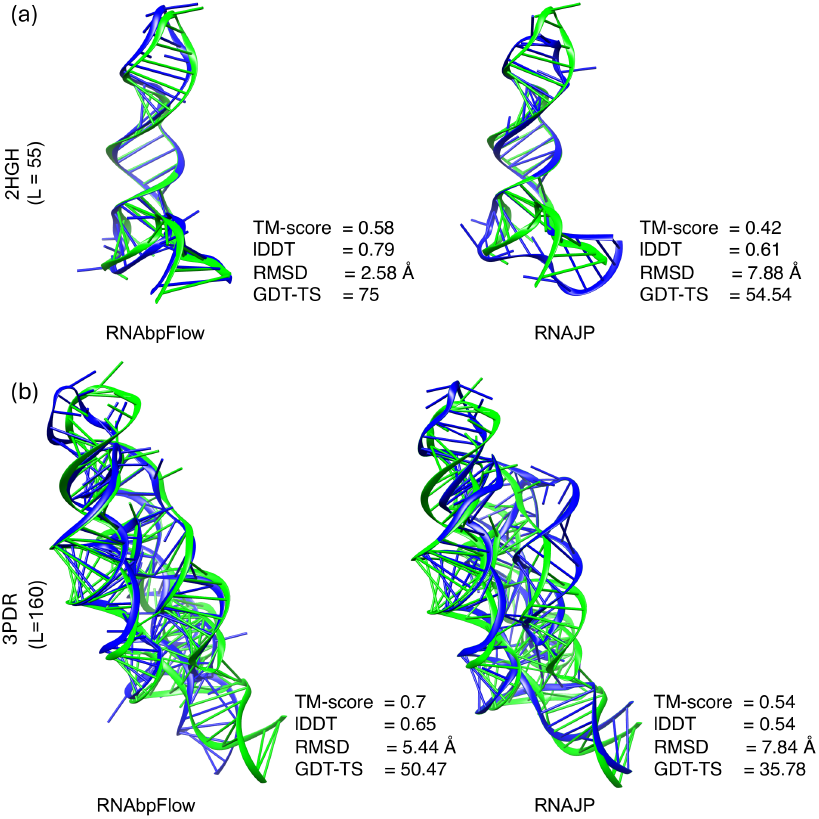
Two representative RNA targets with PDB IDs (a) 2HGH and (b) 3PDR, both containing three-way junctions shown with the predicted structural models colored in blue superimposed on the experimental structures in green and the corresponding evaluation metrics displayed on the bottom right side each superposition.

**Figure 4.**
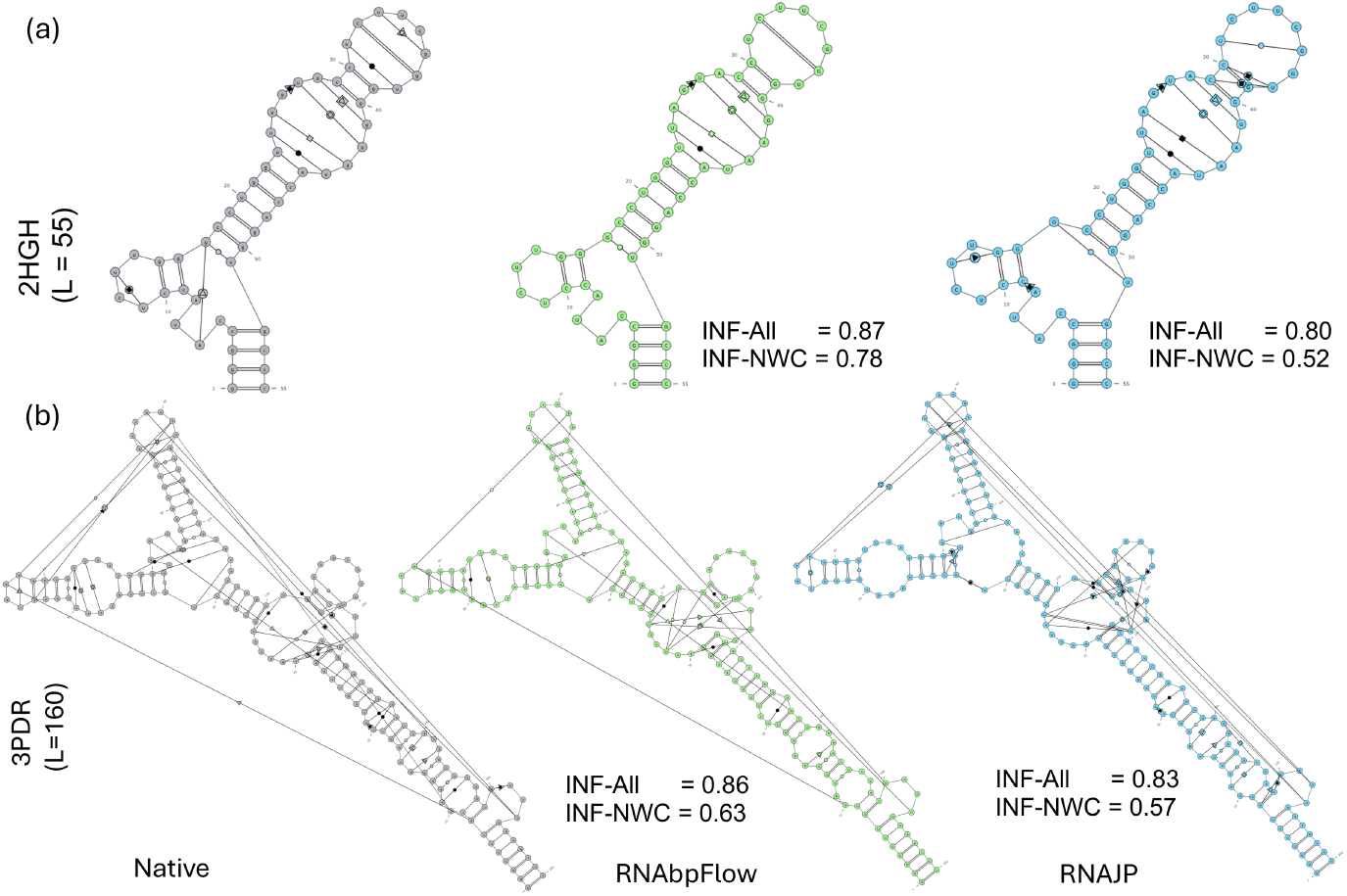
Canonical and non-canonical base pair annotations extracted using RNApdbee 2.0 [44] from the experimental and predicted 3D structures of the two case study targets with PDB IDs (a) 2HGH and (b) 3PDR. Base pair fidelity for the prediction is evaluated and annotated in terms of INF-All (all interactions) and INF-NWC (non-canonical interactions).

For the second case study target 3PDR, a 160-nucleotide target and the largest in the set containing two three-way junctions, our method outperforms RNAJP by achieving an all-atom RMSD of 5.44 Å and a TM-score of 0.7, primarily due to the better packing of the helices and both of the three-way junction regions which is visually evident. By contrast, the best prediction by RNAJP results in an RMSD of 7.84 Å, primarily due to the deviations in one of the three-way junction regions, as shown in **Figure 3**. The extracted base pairs annotated in **Figure 4(b)**, further highlight the trend that RNAbpFlow more accurately captures non-canonical and long-range loop-loop interactions, with fewer false-positives compared to RNAJP.

### 2.5 Performance on CASP15 targets

#### 2.5.1 Sampling performance

**Figure 5** presents a comparison of the sampling performance of RNAbpFlow on four natural CASP15 RNA targets, conditioned on both native and noisy (predicted) base pairs as input. When provided with native base pairs, RNAbpFlow achieves an impressive average over the maximum per-target TM-score of 0.62 and successfully generates at least one 3D structure with a TM-score greater than 0.45 for 100% of the cases. By contrast, when noisy predicted base pairs are used, the performance declines significantly, with the average over the per-target maximum TM-score dropping to 0.48. Despite this performance drop, RNAbpFlow still demonstrates its robust sampling capabilities by generating at least one accurate global fold (TM-score *>* 0.45) for 3 out of 4 targets, showcasing its resilience to noisy 2D information. However, the average sampling performance declines when the input consists entirely of predicted base pair maps, with the average over the mean TM-score per target dropping from 0.42 to 0.32. Such a performance decline highlights the importance of accurate base pairing information in guiding the sample generation process. Additionally, the total number of unique non-canonical base pairs across the three maps for the four targets decreases sharply from 26 to only 3 (with no non-canonical pairs present in 3 out of the 4 targets) when predicted base pairs are used, which could contribute to the observed drop in performance.

**Figure 5.**
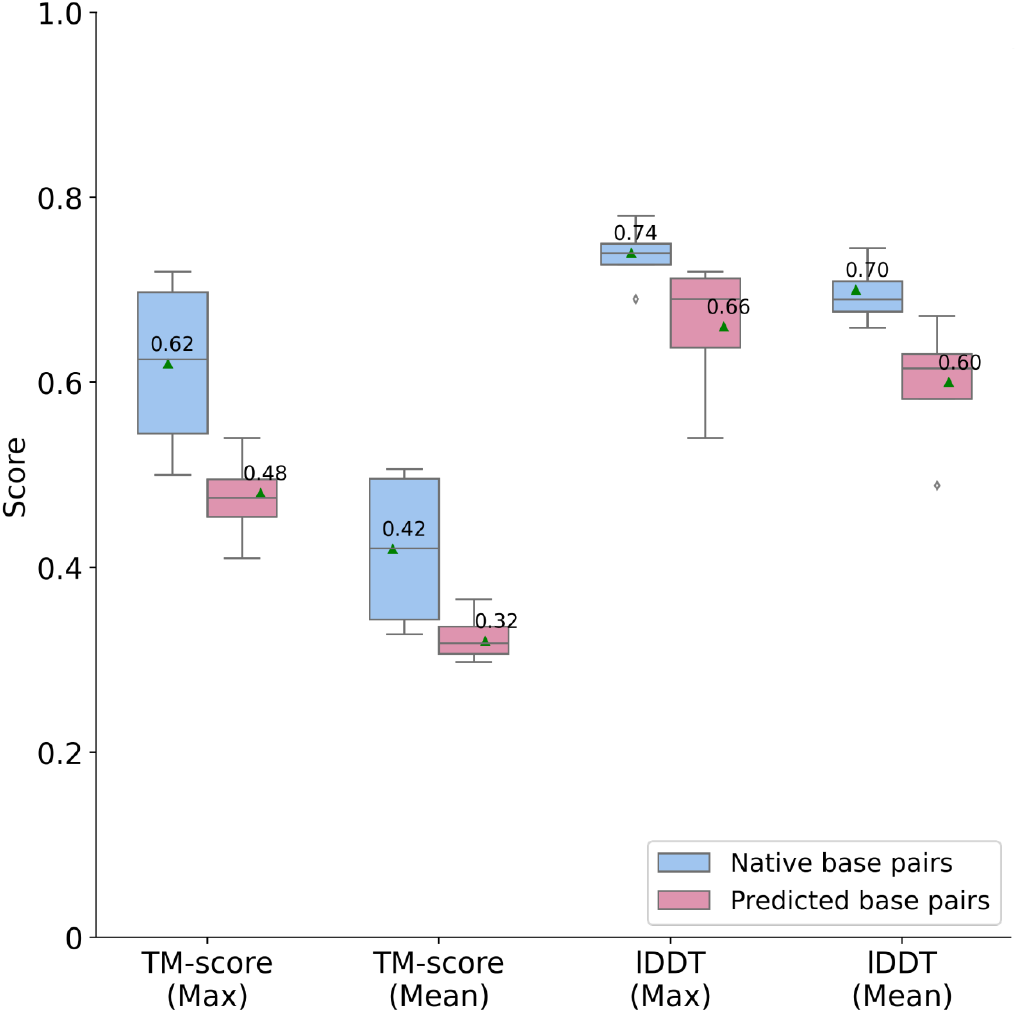
Distributions of the maximum and mean of 1.000 3D structural samples per target in terms of TM-score and lDDT across 4 CASP15 test targets for both native and predicted base pair conditioning.

#### 2.5.2 Predictive modeling performance

To directly compare our conditional flow matching-based RNA 3D structure generation method RNAbpFlow against the state-of-the-art sequence- and base-pair-conditioned RNA 3D structure prediction approaches, we leverage our recently published RNA 3D structure scoring method lociPARSE [45], to select the top structure from the RNAbpFlow generated structural ensemble based on the estimated lDDT score (pMoL). The detailed performance comparison is summarized in **Table 2**, showcasing the ability of RNAbpFlow to predict RNA 3D structures across a wide range of evaluation metrics. RNAbpFlow consistently shows better accuracies than the state-of-the-art deep learning-based method DRfold, achieving a well rounded performance across all the metrics and consistently outperforms all physics- and/or knowledge-based methods across almost all metrics except for clash score where it falls short of the physics-based method Vfold-Pipeline, which employs a post-refinement strategy specifically designed to resolve structural clashes. However, the overall predictive performance of RNAbpFlow can be improved, as none of the 3 accurate global folds generated during sampling is selected as the top structure, highlighting the need for further improvement of the scoring function to identify accurate 3D structure in the ensemble.

**Table 2:**
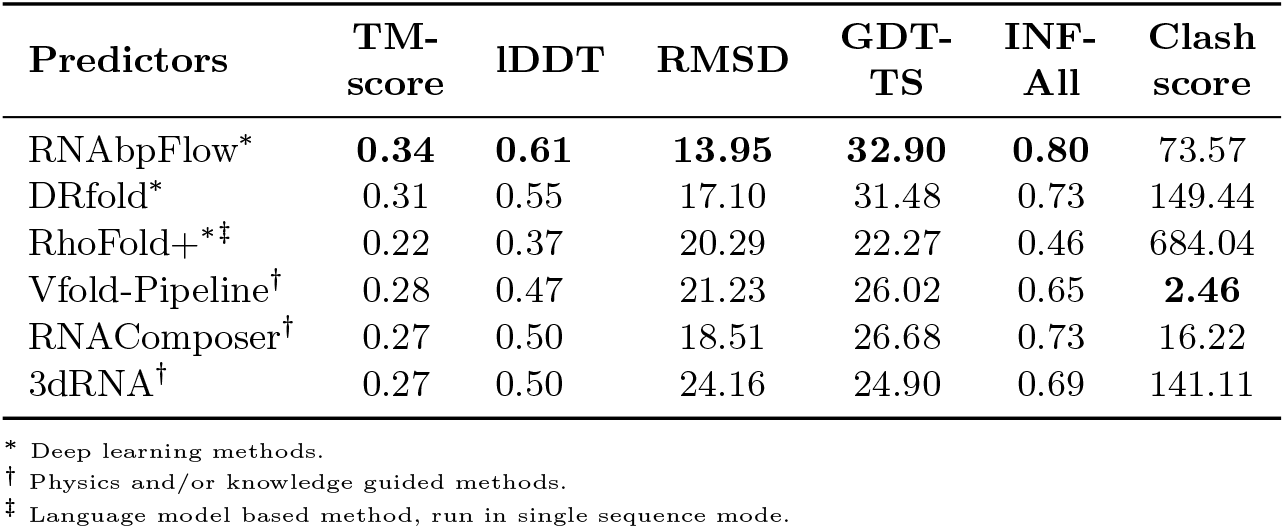
Benchmarking the predictive modeling performance on four natural RNAs from CASP15 with predicted base pairs. Values in bold indicate the best performance.

#### 2.5.3 Impact of native base pairing information

**Table 3** shows how the performance of our method RNAbpFlow varies in the presence of accurate native base pairing information compared to all other competing methods. The predictive modeling performance of our method RNAbpFlow significantly improves in the presence of accurate base pairs, achieving an average TM-score of 0.51 and average RMSD of 7.79, compared to 0.34 and 13.95 TM-score and RMSD respectively, when predicted base pairs are used, showing a 50% improvement in TM-score and a 44.1% reduction in RMSD, which is the highest performance gain compared to all other methods. This highlights the versatility of RNAbpFlow in its ability to achieve a higher performance ceiling by effectively leveraging accurate base pairing information as a key condition in deep generative modeling.

**Table 3:** Comparison of RNAbpFlow with other predictors on CASP15 targets with native base pairing information. Values in bold indicate the best performance.

| Predictors | TM-score | IDDT | RMSD | GDT-TS | INF-All | Clash score |
| --- | --- | --- | --- | --- | --- | --- |
| RNAbpFlow | **0.51** | **0.72** | **7.79** | **49.66** | **0.89** | 46.97 |
| DRfold | 0.36 | 0.67 | 14.47 | 36.82 | 0.85 | 131.68 |
| RNAComposer | 0.32 | 0.59 | 16.81 | 31.01 | 0.84 | 20.73 |
| 3dRNA | 0.30 | 0.54 | 17.90 | 28.08 | 0.72 | 144.51 |
| VFold | 0.28 | 0.58 | 19.69 | 26.44 | 0.80 | **0.64** |

### 2.6 Performance on CASP16 targets

#### 2.6.1 Performance comparison with state-of-the-art methods

AlphaFold 3 (AF3) [28] has emerged as a unified framework for deep learning–based modeling of biomolecular 3D structures including RNA 3D structure prediction, in addition to recent AlphaFold2-inspired architectures such as trRosettaRNA2 [19] and NuFold [22]. In the recently concluded CASP16 challenge, the two automated server methods officially ranked in the top 10 in RNA structure prediction category, AlphaFold 3 (AF3-server) and trRosettaRNA2 (Yang-Server), have demonstrated state-of-the-art performance, alongside top-ranked human expert predictors [46, 47]. Multiple participating groups in CASP16, both human and automated, incorporated AF3 predictions to generate RNA structural ensemble [48]. However, the accuracies of these state-of-the-art methods still remain heavily reliant on the availability of evolutionary information in the form of homologous structural templates and/or deep MSAs [48]. To evaluate the predictive modeling accuracy of our MSA- and template-free method RNAbpFlow on CASP16 targets, we use a training set from the full RNA3DB dataset [34], which is curated from the PDB on 2024-04-26 prior to the start of the CASP16 challenge and obtain 912 RNA sequences of length *<* 200. Following recent literature [49], we further perform data augmentation using a cross-distillation set of additional 2,170 sequences collected from the bpRNA-1m (90) dataset [35], along with their corresponding experimental base pair information. The details of the curation of the cross-distillation set is available in section 4.4. For training, we use base pairs extracted from experimental 3D structures (for RNA3DB) or experimental base pair annotations (for bpRNA-1m (90)) as ground truth. Since RNAbpFlow is trained on RNA sequences up to 200 in length due to the limitation of computational resources, its applicability to larger RNAs (≥200 nucleotides) may be hindered. We, therefore, use 14 publicly available CASP16 targets whose sequence lengths do not exceed 200 nucleotides for benchmarking. During inference, we incorporate predicted base pairs from 15 different prediction methods (based only on sequence information), including those that can handle pseudoknot predictions, to capture a variety of base pairing patterns. The complete lists of 14 publicly available CASP16 targets and the associated base pair prediction methods are provided in **Supplementary Tables S1 and S3**, respectively.

As reported in **Table 4**, our method RNAbpFlow without using any MSA information leads to better accuracy in terms of average of the maximum TM-score and lDDT among the models present in the generated structural ensemble than the two top-performing automated CASP16 servers achieve in their best of five CASP16 submissions despite using MSA and/or template information. To analyze the dependency on MSA and template information of these methods, we compute the target-wise TM-score difference between the best of five CASP16 submissions from either of the methods and the maximum TM-score achieved by the structural ensemble generated by RNAbpFlow. As shown in **Figure 6**, for hard targets (TM-score *<* 0.45) [48] with weak evolutionary signal in terms of shallow MSA depth (Neff 1 ≤ 30) [48], RNAbpFlow consistently yields higher TM-scores in the structural ensemble than both AF3 and trRosettaRNA2, demonstrating the effectiveness of base pair-conditioned structure modeling when evolutionary information is scarce. By contrast, for easy targets such as R1263 or R1264, both AF3 and trRosettaRNA2 outperform RNAbpFlow due to the availability of deep MSA, revealing their dependence on MSA information. Since the search for homologous sequences to obtain sufficiently deep MSA is a challenging task for RNA [2 4, 50] (only 4 out of 14 CASP16 targets have sufficiently deep MSA with Neff *>* 130), our method RNAbpFlow can be an attractive alternative for RNA 3D structure generation conditioned only on base pair information, which is much more conserved across species than sequences [50]. Detailed target-wise results for the two CASP16 servers are available in **Supplementary Table S5**. To benchmark the sampling performance of RNAbpFlow against the state-of-the-art methods in the absence of MSA, we compare our method against the stand-alone versions of AlphaFold 3 (docker version) and NuFold, both run via local installation. We are unable to include trRosettaRNA2 since a stand-alone version of trRosettaRNA2 is not available. For each method, we generate 1,000 structures per target by varying the random seed and running in single-sequence mode by providing only the RNA sequence as input (excluding MSAs). As reported in **Table 4**, the performance of RNAbpFlow is better than AF3 in terms of maximum TM-score and comparable to AF3 in terms of maximum lDDT over the average of 14 targets. At least one correct fold (TM-score *>* 0.45) is present in the generated ensemble of RNAbpFlow for 12 out of 14 targets (85.71%) compared to 8 out of 14 targets (57.13%) for AF3, demonstrating the consistency of RNAbpFlow. Meanwhile, RNAbpFlow yields much better performance compared to NuFold both in terms of TM-score and lDDT. Target-wise results are provided in **Supplementary Table S6** for AF3 and NuFold, and in **Supplementary Table S7** for RNAbpFlow.

**Table 4:**
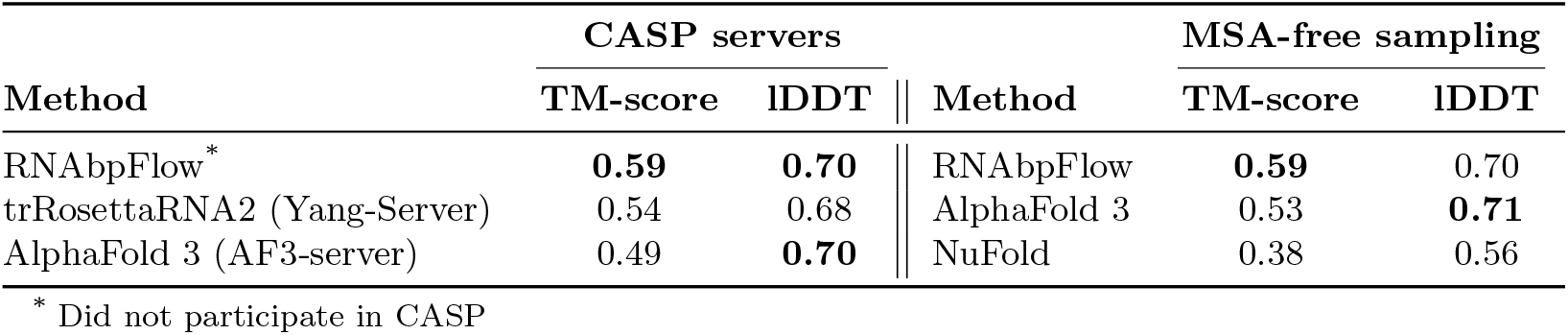
Performance comparison of RNAbpFlow (trained on cross-distillation augmented dataset and predicted base pairs as input during inference) against state-of-the-art methods on CASP16 test set in terms of average of maximum TM-score and lDDT present in the predicted ensemble of each method.

**Figure 6.**
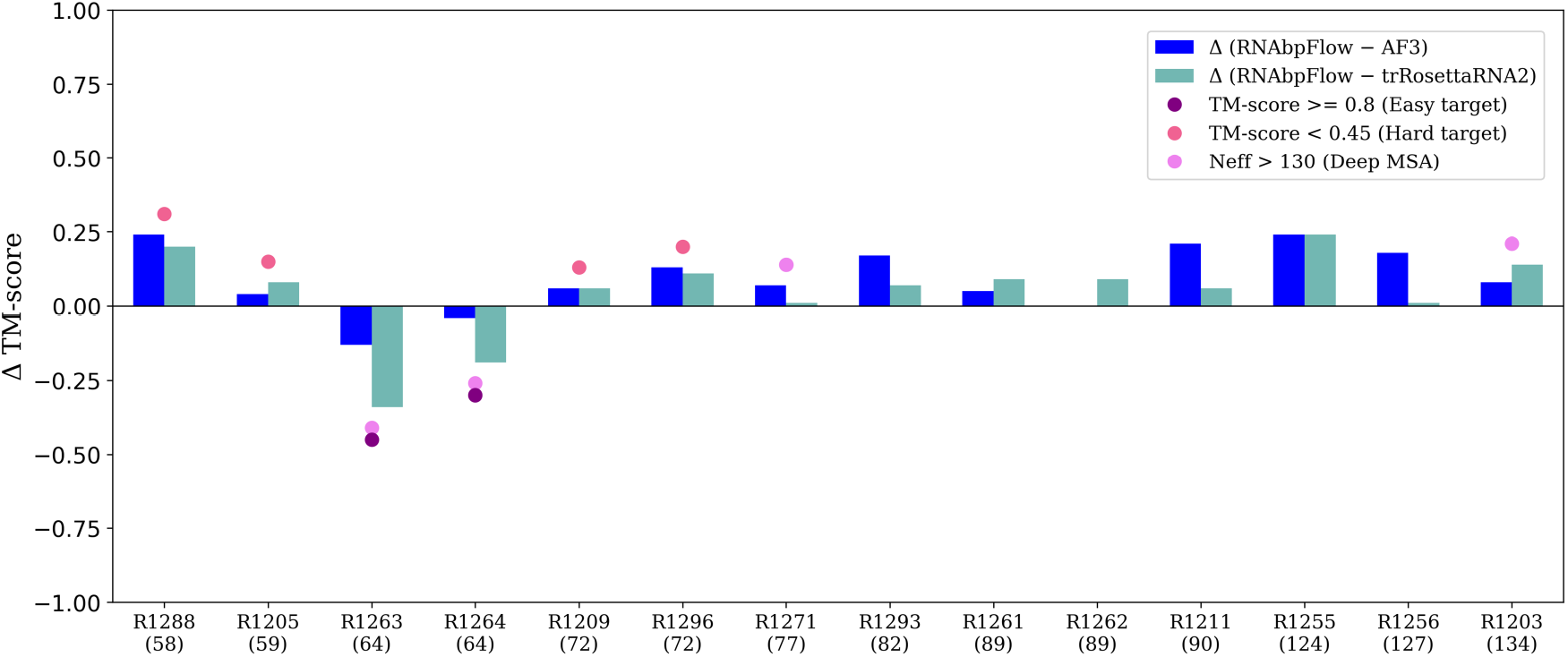
Target-wise TM-score differences between the best structure (maximum TM-score) in RNAbpFlow ensemble (with cross-distillation + predicted base pair) and the best of five submissions from AF3-server (Group 304) and Yang-Server (Group 052) on CASP16 test set sorted by ascending order of the sequence lengths. Positive values indicate targets where RNAbpFlow outperforms the other two methods, highlighting its strength on hard targets and/or sequences with shallow MSAs.

**Figure 7** presents two CASP16 targets with their corresponding best structural models chosen from the generated ensemble for further analysis and comparison with the two top-performing automated CASP16 servers AF3 and trRosettaRNA2. Target R1288 (evaluated nucleotides 1 − 51 due to sequence mismatch) shown in **Figure 7(a)** is an *in vitro* ribozyme having no MSA or template information. It is interesting to note that none of the submissions from AF3 or trRosettaRNA2 in CASP16 result in the correct fold, achieving a maximum TM-score of 0.29 by AF3 (R1288TS304 5) and a TM-score of 0.33 by trRosettaRNA2 (R1288TS052 5), highlighting their limitations in the absence of evolutionary information. Our method RNAbpFlow, on the other hand, successfully recovers the correct fold by attaining a much better maximum TM-score of 0.53. Target R1255 (SL5 SARS-CoV-2) shown in **Figure 7(b)** is one of the several conserved RNA structural elements in the 5^*′*^-UTR of SARS-CoV-2. This stem loop contains a compact 4-way rotated junction at the core, crucial for forming complex 3D folds and drug targeting. While the best sampled structure by RNAbpFlow using predicted base pair inputs correctly identified the orientation of the four helices around the junction, the best submitted structure by both AF3 (R1255TS304 3) and trRosettaRNA2 (R1255TS052 4) contains an expanded junction with two misaligned helices out of four, resulting in 13.25 Å and 10.69 Å all-atom RMSD, respectively, compared to RNAbpFlow’s 4.65 Å. Focusing specifically on the junction loop region defined by eight nucleotides (24, 25, 69, 70, 94, 95, 104, 105), RNAbpFlow achieves a junction RMSD of 2.41 Å, whereas AF3 reaches 4.1 Å. This highlights the importance of accurate junction-loop modeling, where even subtle misorientation in local motifs can drastically affect the overall fold, as reflected in the all-atom RMSD differences observed in this case study.

**Figure 7.**
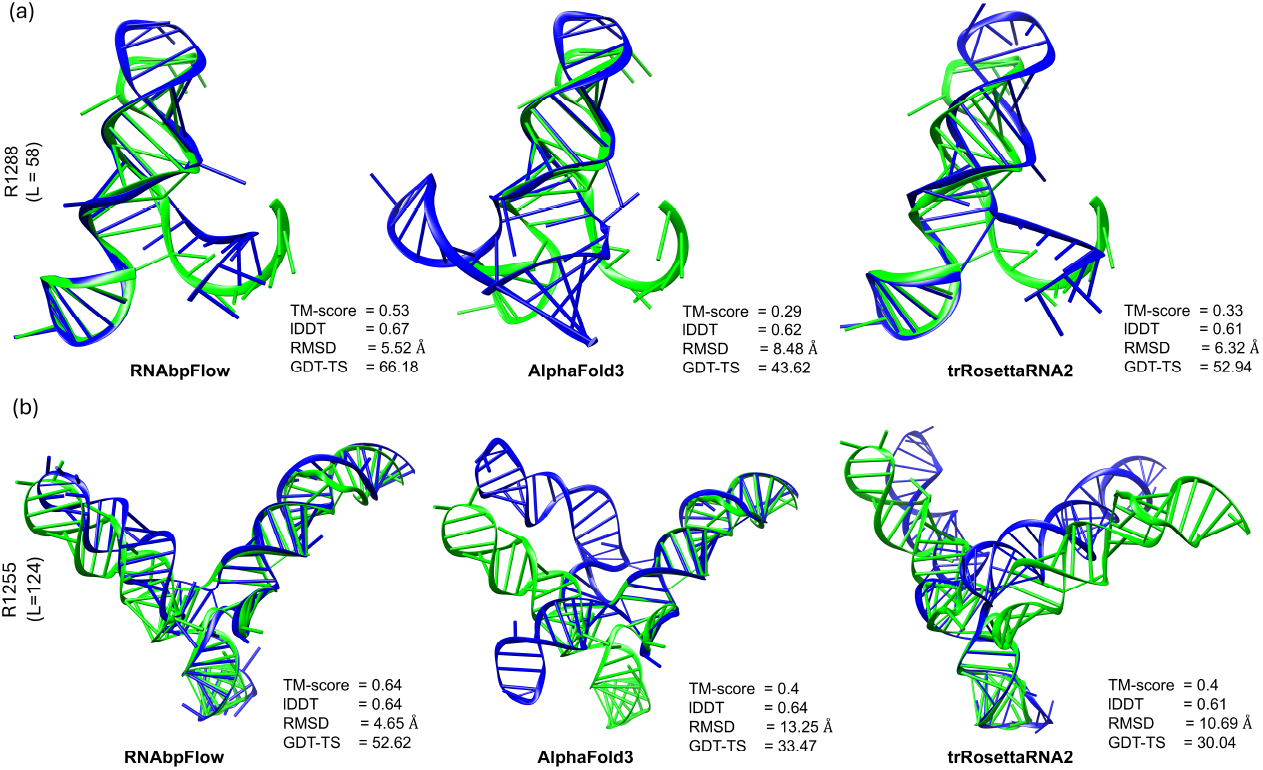
Two representative targets from CASP16 set (a) R1288 and (b) R1255 with the predicted structural models colored in blue superimposed on the experimental structures in green and the corresponding evaluation metrics displayed on the bottom right sides of the structural superpositions.

#### 2.6.3 Contribution of data augmentation and impact of base pair accuracy

To evaluate the contribution of data augmentation via cross-distillation training of RNAbpFlow and the impact of base pair accuracy, we generate 1,000 3D structural models for each CASP16 target using both native and predicted base pairs as inputs with and without distillation. As reported in **Table 5**, data augmentation via cross-distillation training yields substantial gains over the “no distillation” variant regardless of the nature of base pair information (i.e., predicted vs. experimental) used for conditional generation (target-wise results with and without distillation are available in **Supplementary Table S7**). With predicted base pair conditioning, the average over the maximum TM-score increases from 0.51 to 0.59, a relative improvement of 15.7%, and the average over the maximum lDDT increases from 0.63 to 0.70 with a relative improvement of 11.1%. With experimental base pair conditioning, the performance improves significantly, attaining an average over the maximum of 0.65 and an average over the maximum lDDT of 0.73. To analyze the fidelity with which the input base pairs are realized by RNAbpFlow predicted structures, we compute the INF score between the input base pairs (native or predicted) and the output base pairs extracted from the resulting 3D structures using RNAView [51]. **Supplementary Table S10** highlights that when native base pairs are provided, RNAbpFlow generates structures with strong agreement to the input with an average INF value of 0.89. Furthermore, when provided with noisy (predicted) base pairing information that differs from the native input, RNAbpFlow still reproduces these incorrect input base pairs with slightly lower but still high fidelity (average INF = 0.82), demonstrating the ability of RNAbpFlow to tightly conform to the provided input base pairs regardless of their accuracy. Overall, the results demonstrate that RNAbpFlow can recover correct 3D structures, both in terms of TM-score and lDDT, when supplied with accurate experimental base pairs; and while an accuracy gap remains when predicted base pairs are used compared to experimental inputs, data augmentation via cross-distillation during training helps to close the gap to some extent. Target-wise metrics using native base pairs as input are provided in **Supplementary Table S4**, while the relative change in performance compared to predicted input base pairs as well as sampling statistics (percentages of correct folds in the ensemble) are shown in **Supplementary Tables S8 and S9**, respectively.

**Table 5:** Sampling performance in terms of maximum and mean scores of 1,000 3D structures for each target in the CASP16 test set based on different variants of training and inference of RNAbpFlow. All models are trained using experimental base pair annotations.

| Method | Training | Inference | TM-score |  | IDDT |  |
| --- | --- | --- | --- | --- | --- | --- |
|  |  |  | Max | Mean | Max | Mean |
| RNAbpFlow | w/ distillation | experimental bp | **0.65** | **0.49** | **0.77** | **0.73** |
| RNAbpFlow | w/ distillation | predicted bp | 0.59 | 0.38 | 0.70 | 0.60 |
| RNAbpFlow | w/o distillation | predicted bp | 0.51 | 0.35 | 0.63 | 0.55 |

### 2.7 Ablation study and hyperparameter selection

To evaluate the importance of various loss components and their contributions to sampling performance, along with the architectural hyperparameters used during RNAbpFlow’s training, we perform ablation study using the RNA3DB dataset described in Section 4.4. **Figure 8** and **Table 6** report the sampling performance in terms of TM-score and lDDT on the RNA3DB test set, calculated as the average over the per-target maximum and mean scores across 52 RNA3DB test targets for different ablated variants. **Figure 8** shows that RNAbpFlow achieves the best performance when all three loss components are used, with a consistent performance decline when any loss component is removed. For instance, excluding either or both 2D and 3D base pair loss components (i.e., *bp*2*D* and/or *bp*3*D*) leads to a noticeable drop in average TM-score and lDDT, underscoring the importance of incorporating base pairs conditioning in the training. Furthermore, the average lDDT achieved using only the SE(3)-flow matching loss is 0.44, significantly lower than the 0.66 average lDDT obtained when incorporating torsion loss and 2D and 3D base pair losses altogether. This demonstrates the effectiveness of all the auxiliary loss terms used in RNAbpFlow.

**Table 6:**
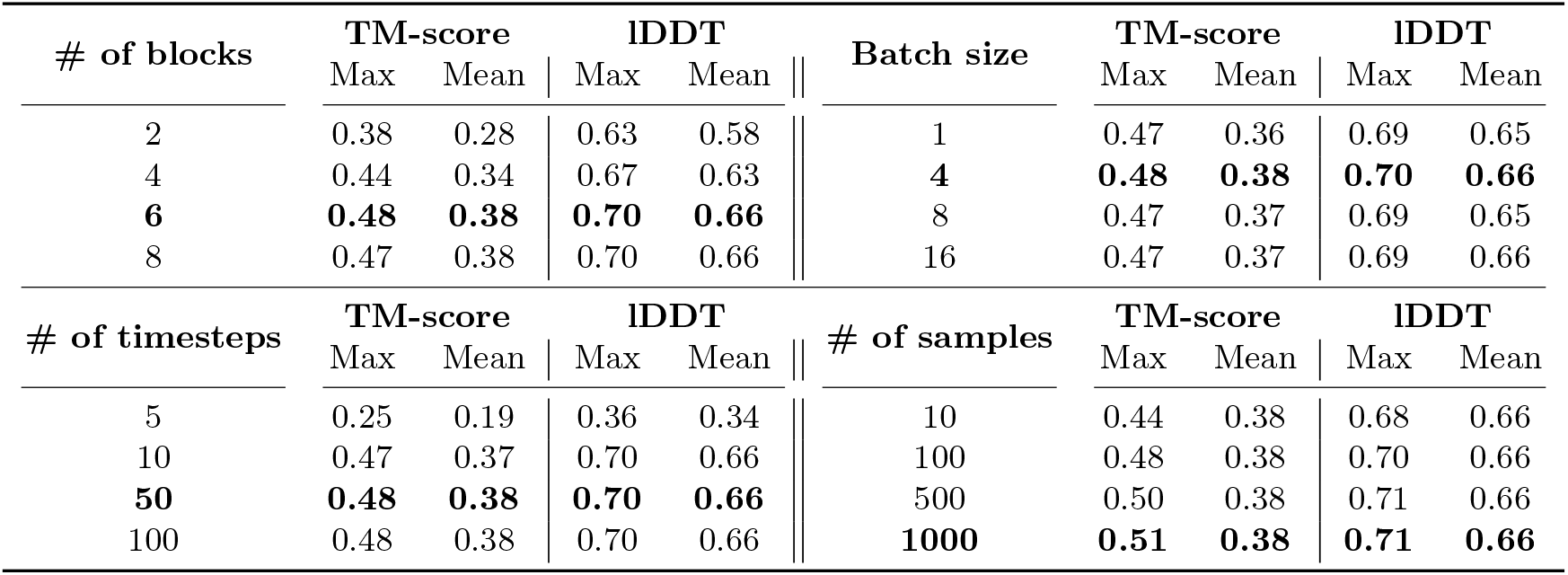
Hyperparameter selection based on number of structure-module blocks, batch size, number of timesteps used to generate a sample RNA 3D structure, and number of samples generated per target on the RNA3DB test set. Values in bold indicate the best performance.

| # of blocks | TM-score |  | IDDT |  | Batch size | TM-score |  | IDDT |  |
| --- | --- | --- | --- | --- | --- | --- | --- | --- | --- |
|  | Max | Mean | Max | Mean |  | Max | Mean | Max | Mean |
| 2 | 0.38 | 0.28 | 0.63 | 0.58 | 1 | 0.47 | 0.36 | 0.69 | 0.65 |
| 4 | 0.44 | 0.34 | 0.67 | 0.63 | **4** | **0.48** | **0.38** | **0.70** | **0.66** |
| **6** | **0.48** | **0.38** | **0.70** | **0.66** | 8 | 0.47 | 0.37 | 0.69 | 0.65 |
| 8 | 0.47 | 0.38 | 0.70 | 0.66 | 16 | 0.47 | 0.37 | 0.69 | 0.66 |
| # of timesteps | TM-score |  | IDDT |  | # of samples | TM-score |  | IDDT |  |
|  | Max | Mean | Max | Mean |  | Max | Mean | Max | Mean |
| 5 | 0.25 | 0.19 | 0.36 | 0.34 | 10 | 0.44 | 0.38 | 0.68 | 0.66 |
| 10 | 0.47 | 0.37 | 0.70 | 0.66 | 100 | 0.48 | 0.38 | 0.70 | 0.66 |
| **50** | **0.48** | **0.38** | **0.70** | **0.66** | 500 | 0.50 | 0.38 | 0.71 | 0.66 |
| 100 | 0.48 | 0.38 | 0.70 | 0.66 | **1000** | **0.51** | **0.38** | **0.71** | **0.66** |

**Figure 8.**
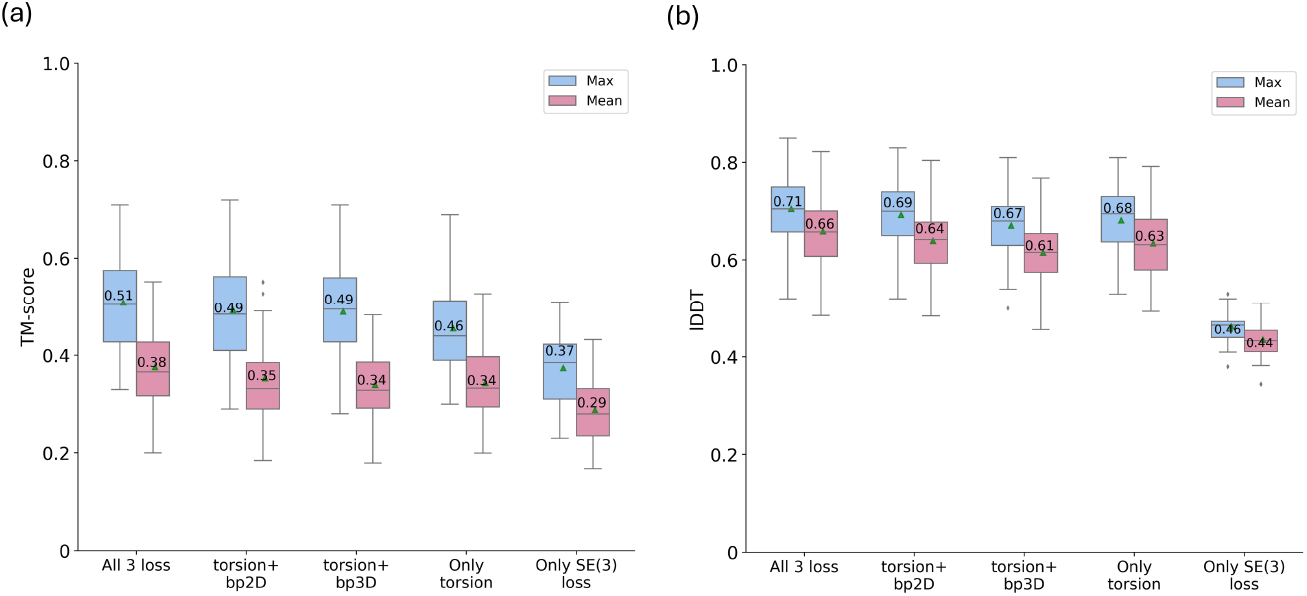
Distributions of the maximum and mean of 1,000 3D structural samples per target in terms of (a) TM-score and (b) lDDT distribution across 52 RNA3DB test targets for five d ifferent lo ss co mbinations. Th e gr een triangles indicate averages.

**Table 6** highlights the impact of hyperparameter selection on sampling performance. The number of structural module blocks significantly influences TM-score and lDDT, with 6 blocks providing the best balance between model complexity and computational cost. Similarly, a batch size of 4 is sufficient for training efficiency without compromising accuracy. The efficiency of the flow matching formulation is also evident from **Table 6**, as our method RNAbpFlow achieves near-optimal results with just 10 timesteps, with marginal gains observed at 50 timesteps, demonstrating the practical advantage of our approach for large-scale RNA 3D structure sampling with minimal computational overhead. Additionally, increasing the number of generated samples consistently improves performance, with 1,000 samples per target yielding the best performance. These findings justify our hyperparameter choices used in this work: 6 structure-module blocks, a batch size of 4, 50 timesteps, and 1,000 samples generated per target.

## 3 Conclusion

In this work, we developed RNAbpFlow, the first sequence- and base-pair-conditioned all-atom R NA 3D structure generation method based on SE(3)-equivariant flow matching model. Experimental results demonstrate that the introduction of base pairing conditioning leads to performance improvements, and the accuracy gain is connected to the quality of the base pairs. Free from the confines of sequence- and structural-level homology, RNAbpFlow enables direct generation of all-atom RNA 3D structural models in an end-to-end manner, thereby opening promising avenues for RNA conformational dynamics through large-scale structural ensemble generation in atomic detail.

## 4 Methods

### 4.1 Model Input

Our method operates independently of any MSA or template information, relying solely on sequence and base pairing (2D) information for conditional generation of RNA 3D structure. Specifically, given an RNA sequence of length *N* as input, we encode the nucleotide sequence using one-hot encoding, represented as a binary vector of 4 entries corresponding to the four types of nucleotides (A, U, C, G). For base pairing information during the training process, we extract 2D annotations from experimental (native) 3D structures using three different software tools: RNAView [51], MC-Annotate [52], and DSSR [53] and represent them as three separate 2D binary maps. However, despite analyzing the same 3D coordinates, each method applies its own geometric and hydrogen-bond cutoffs (e.g., which edges interact, planarity/angle limits, minimum H-bonds) which can result in non-identical base pair annotations depending on these subtle interpretation and small geometrical variations [41] (**Supplementary Table S11**). To capture these diverse canonical and non-canonical base pairing information, we provide all three maps as input for the bias term in our denoiser architecture as edge features. During the sampling of CASP15 and RNA3DB test targets, in the absence of native base pairing information, we use sequence-based predicted base pairs from three different RNA 2D structure predictors, namely MXfold2 [54], SimFold [55], and ContextFold [56]. For the CASP16 targets, in the absence of native base pairing information, we use a diverse set of 15 different RNA 2D structure predictors, a list of which is provided in **Supplementary Table S3**.

### 4.2 Network Architecture

#### 4.2.1 RNA Frame representation

We represent each nucleotide in a geometric abstraction using the concept of rigid body frames. Each nucleotide frame in the form of a tuple is defined as a Euclidean transform *T* = (*r, x*), where *r* ∈*SO*(3) is a rotation matrix and *x* ∈ ℝ^3^ is the translation vector that can be applied to transform a position in local coordinates to a position in global coordinates.

#### 4.2.2 SE(3) flow matching on frames

Flow matching (FM) [57] is a class of deep generative models in which the goal is to learn a velocity field (or flow) that matches the probability flow of the data distribution to transform a simple distribution such as Gaussian to the desired complex data distribution in high-dimensional space. Flow matching directly learns this velocity field that describes how points should move from the simple distribution to the target distribution without completely destroying the data distribution. By integrating an ordinary differential equation (ODE) over this learned vector field, FM offers simpler trajectories towards achieving target distribution offering huge computational speed-up for large-scale sample generation.

In the context of generative modeling, the geodesic path describes a smooth transformation of one probability distribution into another while minimizing distortion. The concept of geodesics generalizes the notion of shortest paths in non-Euclidean spaces, enabling efficient computation and interpretation. Given a noisy frame *T*_0_ sampled from a simple prior density *p*_0_(*T*_0_) and the experimental frame *T*_1_ sampled from a target distribution *p*_1_(*T*_1_), the geodesic path connecting two points **T**_0_ and **T**_1_ in a combination of *simple manifolds* such as ℝ^3^ and *SO*(3) can be expressed using exponential and logarithmic maps following the generalization of flow matching to Riemannian manifolds [58] in the following way:

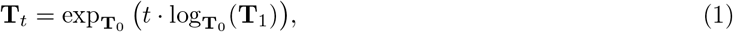

where 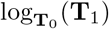 is a vector in the tangent space of **T**_0_ pointing toward **T**_1_, and *t* ∈ [0, 1] parameterizes the sequence of probability distributions a.k.a *probability path p*_*t*_ between the two distributions *p*_0_ and *p*_1_. The conditional flow *T*_*t*_ constructed in Eq. 1 can be decomposed into separate individual flows for the simplification of training procedure in the following way:

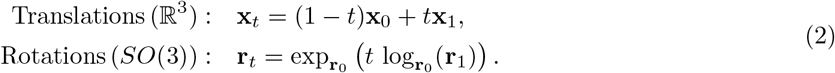

where the prior distribution *p*_0_(*T*_0_) during training takes the form of *p*_0_(*T*_0_) = *IGSO*3(*σ* = 1.5) ⊗ 𝒩 (0, *I*_3_) where random translation *x*_0_ is sampled from the unit Gaussian distribution 𝒩(0, *I*_3_) and random rotation is sampled from *IGSO*3(*σ* = 1.5) following [30, 31, 59] for better performance. During training, the parameter *t* is sampled from 𝒰([0, 1 − *ϵ*]) where *ϵ* = 0.1 is chosen for training stability.

#### 4.2.3 Denoiser model

The objective of our flow matching method is to learn the parametrized vector field **u**_*t*_, which represents a smooth, time-dependent (*t*) map that generates an ordinary differential equation (ODE) that describes the transformation between two distributions: *p*_0_(*T*_0_) (noisy frames) and *p*_1_(*T*_1_) (ground truth frames). To learn this mapping from noisy samples to cleaner ones, we train a parameterized denoiser model **v**_*θ*_(**T**_*t*_, *t*) which will predict clean frames given corrupted ground truth frames **T**_*t*_ at time *t*. Following FrameFlow [31], we use the structure module from AlphaFold2 [17] as the neural architecture of our denoiser model.

#### 4.2.4 Sampling strategy

During conditional sampling of RNA 3D structures, our framework takes a random initialization of the backbone frames *T* = (*r, x*), where translation *x* is sampled from a unit Gaussian distribution 𝒩 (0, *I*_3_) in ℝ^3^ and rotation *r* is sampled from a uniform distribution in SO(3). During inference, instead of the linear scheduler for rotation matrices, we use the exponential scheduler *e*^−*ct*^ with *c* = 10 for better performance. Thus, our SO(3) flow for rotation in Eq. 2 changes according to the following equation:

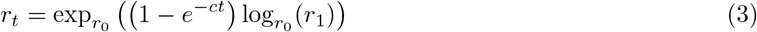

Based on the specified number of timesteps, we choose a set of values for *t* where *t* ∼ *𝒰* ([0, 1]). Starting from the random set of frames, we iteratively update the frame representations using the predictions from previous steps using our learned denoiser model with the specified condition at each timestep *t* in the following way:

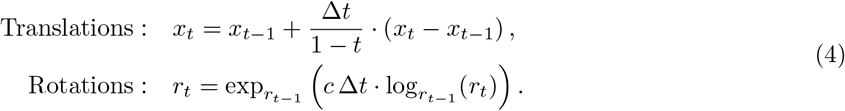

### 4.3 Training of RNAbpFlow

Our training objective contains multiple loss function components related to base pair conditions and allatom structure generation. The primary loss function for this framework is the same SE(3) loss formulated in FrameFlow [31] namely, the vector field loss in SE(3). To train our denoiser model **v**_*θ*_, two separate components of this loss are calculated for the predicted rotation 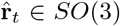 *SO*(3) and translation 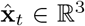 given the corrupted frames **T**_*t*_ at time *t* as shown below.

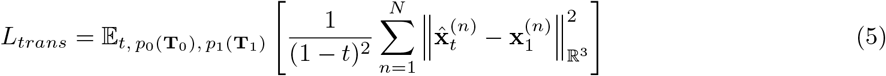

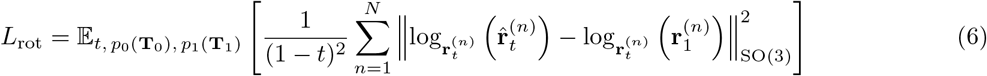

where *N* is the total number of frames. To predict the 9 torsion angles 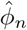, we use an additional head and calculate the torsional loss in the following way by comparing with the experimental angles *ϕ*:

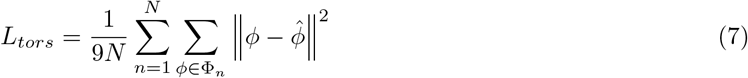

Our base pair-augmented auxiliary loss function components are described below. At the 3D level, we optimize the predicted distance between the annotated base pairs (*m, n*) extracted from all three base pair annotation methods, termed the *bp*3*D* loss, as follows:

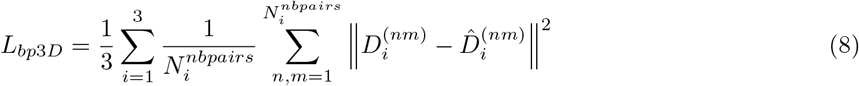

where 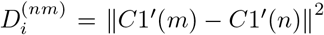 denotes the actual distance between the *C*1^*′*^ atoms of the nucleotide pair (*m, n*) denoted by the *i*^*th*^ 2D structure annotation method and 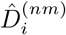 is the predicted distance. At the 2D level, we utilize an additional head to reduce the dimensionality of the predicted pair features (*ŜS*) and compute the BCEWithLogitsLoss against the three experimental 2D maps (*SS*), termed the *bp*2*D* loss, as described below:

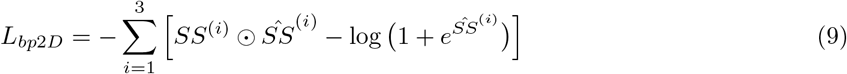

Our final combined weighted loss function is shown below:

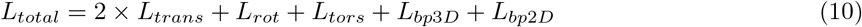

We train our model in PyTorch-Lightning using the Adam optimizer with a learning rate of 0.0001. The distributed training process runs on 8 NVIDIA H100 GPUs for 1000 epochs, taking approximately 25 hours to train on the augmented training set via cross-distillation.

### 4.4 Training and benchmark datasets

To develop our method RNAbpFlow, we utilize two separate training sets and corresponding non-redundant benchmark sets. For the primary method development and internal benchmarking, we use the recently published RNA3DB dataset [34], which is both sequentially and structurally non-redundant making it highly suitable for training and benchmarking deep learning models. We use the version of RNA3DB parsed from the PDB [12] on April 26, 2024 and select a representative set of RNA sequences from the original train-test split they provide for our purposes. To ensure that the dataset contains only high-quality native structures, we apply several filtering steps such as excluding structures with only one atom per nucleotide, removing protein residues from RNA structures, extracting contiguous sequences from experimental structures to address mismatches between the provided FASTA sequences and the experimental 3D structures for preserving base pairing integrity, and excluding sequences lacking any base pairs in their corresponding native structures. Finally, we only retain sequences with a minimum length of 30 and a maximum length of 200 to ensure efficient training. This results in a clean training set consisting of 573 RNA sequences (excluding component #0, which comprises of sequences with no hit to any Rfam family at an E-value threshold of 1.0, reducing the chance of data leakage) and 52 test sequences for benchmarking our method development. Both training and test sets originate from RNA3DB and follow the non-redundant splitting strategy defined in the original publication [34], which ensures no sequence or structural overlap between the training and test sets.

To compare RNAbpFlow against RNAJP [33], a recent MD simulation-based method for RNA 3D structure sampling based on given 2D structure, we use the dataset of 22 RNAs containing three-way junctions as used in the original RNAJP study. We exclude multimeter structures from the dataset and apply CD-HIT-est-2D between the remaining sequences and our training set of 573 RNA sequences from RNA3DB to avoid redundancy. This reduces the benchmark set to 12 RNAs (**Supplementary Table S2**), for which we download the predicted all-atom decoy structures generated by RNAJP from their publicly available repository at https://rna.physics.missouri.edu/RNAJP/index.html.

To benchmark on CASP15 targets, we do not reuse the RNA3DB training set anymore. Instead, we construct a completely new and independent training set sourced from trRosettaRNA [14] using the same rigorous filtering steps as above. This training set only includes RNA chains released in the Protein Data Bank (PDB) prior to January 2022 [14]. Since the release date of the first target in the CASP15 challenge was May 2022, this guarantees that the CASP15 test targets were published more recently than any targets available in the training set. To ensure further sequence-level non-redundancy, we apply MMseqs2 clustering [60] with a 20% identity threshold between CASP15 RNAs and this training set, which results in a training set of 874 sequences with no sequence-level redundancy with CASP15 targets. Four natural CASP15 targets (R1107, R1108, R1149, R1156) are selected for evaluation, each confirmed to be free of chain breaks in its corresponding experimental structure deposited in PDB to preserve base pairing consistency.

To benchmark the performance of RNAbpFlow on the CASP16 targets, we merge all components provided by RNA3DB into a single training set to train RNAbpFlow. The first CASP16 target was released in May 2024, whereas the RNA3DB training set contains PDBs released on or before April 26, 2024, thus emulating a strictly blind experiment as followed by other participating groups in the CASP16 challenge. Following the filtering procedure described above, we obtain a total of 912 RNA sequences (maximum length of 200) and their corresponding experimental 3D structures. For the benchmark set, we collect the 14 targets of length *<* 200 currently publicly available on the official CASP16 website, a list of which is available in **Supplementary Table S1**. We additionally use a cross-distillation set from bpRNA-1m (90) [35] to investigate the effect of data augmentation. To this end, we first obtain the bpRNA-1m(90) set [35], which is comprised of 28,370 sequences (and their corresponding secondary structures), as originally curated by the authors after removing any sequences sharing *>* 90% identity over at least 70% of their length. We then filter this set by sequence length to retain only those sequences between 30 and 200 nucleotides. To further reduce redundancy, we apply MMseqs2 clustering [60] with a 20% identity threshold, yielding 11,048 representative sequences. Next, we remove any of these 11,048 sequences that share *>* 20% identity with the combined RNA3DB set of 912 training sequences and all CASP16 targets, again using MMseqs2, resulting in 10,107 non-redundant sequences. From this pool, we exclude sequences whose bpRNA-1m(90) provided secondary structures contain *<* 10 base-pairs, reducing the set to 9,289 sequences. For data augmentation via cross-distillation during training, we predict structures for these 9,289 sequences using AlphaFold 3 [28] (Docker version) following recent literature [49]. We do not use any MSA information during AlphaFold 3 prediction to generate the cross-distillation dataset for augmentation to ensure that the training of our method RNAbpFlow remains free from the influence of any evolutionary information. To maximize the estimated confidence scores of the AlphaFold 3 predicted structures in the augmented training set, we filter the predictions using the self-assessment confidence measures plDDT (predicted local distance different test) and PAE (predicted aligned error) of AlphaFold 3. Specifically, we only retain the predictions when plDDT ≥ 60 and PAE ≤ 15, resulting in a final cross-distillation set of 2,170 high-confidence RNA structures. Overall, the training set comprises of 912 experimental PDB structures and 2,170 high-confidence predicted structural models for data augmentation via cross-distillation (for a combined total of 3,082), maintaining an approximate 1:3 ratio of PDB to cross-distillation data within each training batch [49].

### 4.5 Performance evaluation and competing methods

To evaluate the performance of our method, RNAbpFlow, compared to other state-of-the-art 3D structure sampling and prediction methods for RNA, we use a wide range of evaluation metrics. To evaluate global fold similarity, we calculate the TM-score (based on C3^*′*^) using US-align [38] and GDT-TS (based on C4^*′*^) using the LGA program for RNA [39]. For the assessment of local environment fitness, we compute lDDT [40] using OpenStructure [61] version 2.8 and the clash score metric using MolProbity package [42]. Furthermore, we utilize the CASP-RNA pipeline [43] to (1) evaluate the full atomic structural accuracy by calculating the all-atom RMSD and (2) compute the INF-All score [41], which quantifies how well a structural model reproduces the base interactions of the reference (usually an experimentally resolved native structure) by considering canonical, non-canonical, and stacking interactions.

Since our method RNAbpFlow relies purely on sequence- and base-pair-conditioned SE(3)-equivariant flow matching model for RNA 3D structure generation without utilizing MSAs or template information, we choose two deep learning-based methods that can only take sequence and base pair information for RNA 3D structure prediction for the sake of a fair performance comparison. These include DRfold that integrates end-to-end and geometric potentials and RhoFold+ which leverages language model. Both DRfold and RhoFold+ are installed and run locally with their default parameters, with RhoFold+ run in single-sequence mode. For the CASP16 targets, we additionally compare RNAbpFlow against state-of-the-art diffusion-based biomolecular prediction method AlphaFold 3 [28] and two other AlphaFold2-inspired RNA 3D structure prediction methods NuFold [22] and trRosettaRNA2 [19]. In addition to these deep learning-based approaches, we compare against three physics- and/or knowledge-based methods: RNAComposer [10], 3dRNA [36], and Vfold-Pipeline [37]. RNAComposer and 3dRNA are accessed via their respective web servers, while Vfold-Pipeline is installed and run locally. For these methods, base pairing information is provided in dot-bracket notation format, as predicted by MXfold2 software. To maintain a fair comparison and accurately assess predictive performance, we do not employ any post-prediction optimization or refinement procedures for structure prediction across all methods, except in the cases of Vfold-Pipeline and RNAComposer, where the option to switch off the post-prediction refinement functionality is not available.

## 5 Acknowledgements

This work was partially supported by the National Institute of General Medical Sciences (R35GM138146 to D.B.) and the National Science Foundation (DBI2208679 to D.B.).

