## Supplementary Information for "RNAbpFlow: Base pair-augmented SE(3)-flow matching for conditional RNA 3D structure generation"

### Supplementary Tables

#### S1 List of targets in CASP16 benchmark set

Table S1: Summary of 14 publicly available RNA targets (length < 200) from CASP16 competition, including effective number of sequences in MSA(Neff), highest template matching score against existing targets in PDB, and difficulty level. Neff and TM-score values are taken from official CASP16 assessment [1]. TM-score < 0.45 is defined as hard target whereas TM-score ≥ 0.8 is considered as easy target.

| ID | Length | Neff | TM-score | Difficulty |
|---|---|---|---|---|
| R1288 | 58 | 19 | 0.384 | Hard |
| R1205 | 59 | 9 | 0.402 | Hard |
| R1263 | 64 | 554 | 0.930 | Easy |
| R1264 | 64 | 554 | 0.793 | Easy |
| R1209 | 72 | 46 | 0.363 | Hard |
| R1296 | 72 | 48 | 0.430 | Hard |
| R1271 | 77 | 1742 | 0.699 | Medium |
| R1293 | 82 | 11 | 0.589 | Medium |
| R1261 | 89 | 14 | 0.730 | Medium |
| R1262 | 89 | 3 | 0.493 | Medium |
| R1211 | 90 | 89 | 0.664 | Medium |
| R1255 | 124 | 14 | 0.730 | Medium |
| R1256 | 127 | 3 | 0.493 | Medium |
| R1203 | 134 | 859 | 0.630 | Medium |

#### S2 List of targets in RNAJP benchmark set

Table S2: List of 12 PDBs used in sampling performance comparison against RNAJP method.

| ID | Length |
|---|---|
| 3PDR_X | 160 |
| 2N1Q_A | 155 |
| 2NBX_A | 108 |
| 2OIU_Q | 71 |
| 2NC1_A | 67 |
| 3R4F_A | 66 |
| 3EGZ_B | 65 |
| 1MMS_C | 58 |
| 1DK1_B | 57 |
| 2HGH_B | 55 |
| 2MHI_A | 53 |
| 3E5C_A | 53 |

#### S3 Secondary structure prediction methods for CASP16 sampling

Table S3: List of 15 secondary structure prediction methods used in CASP16 sampling. Since RNAbpFlow can incorporate 3 base pair maps at the same time, we generate 200 3D structures for each set of 3 methods, totaling 1000 structures per target.

| Method | Pseudoknot Available | Deep Learning |
|---|---|---|
| SPOT-RNA [2] | ✓ | ✓ |
| UFold [3] | ✓ | ✓ |
| KnotFold [4] | ✓ | ✓ |
| MXfold2 [5] | ✗ | ✓ |
| Iterative HFold [6] | ✓ | ✗ |
| HotKnot v0.2 [7] | ✓ | ✗ |
| RNAstructure [8] | ✓ | ✗ |
| IPKnot [9] | ✓ | ✗ |
| ContextFold [10] | ✗ | ✗ |
| CONTRAfold [11] | ✗ | ✗ |
| EternaFold [12] | ✗ | ✗ |
| LinearFold [13] | ✗ | ✗ |
| PETfold [14] | ✗ | ✗ |
| RNAfold [15] | ✗ | ✗ |
| SimFold [16] | ✗ | ✗ |

#### S4 Target-wise performance of RNAbpFlow on CASP16 with native base pairs

Table S4: Per-target TM-score and lDDT statistics for RNAbpFlow inference on 14 CASP16 targets. The columns represent maximum and average over 1000 3D structures generated for each target with base pair conditions extracted from the corresponding experimental 3D structures available in PDB. Bold values denote the presence of atleast one correct fold in the ensemble (TM-score > 0.45 or lDDT > 0.75.)

| **ID** | **TM-score** | | **lDDT** | |
|---|---|---|---|---|
| | Max | Mean | Max | Mean |
| R1288 | 0.43 | 0.35 | **0.78** | 0.74 |
| R1205 | **0.47** | 0.36 | 0.69 | 0.63 |
| R1263 | **0.77** | 0.62 | **0.87** | 0.82 |
| R1264 | **0.74** | 0.59 | **0.85** | 0.80 |
| R1209 | 0.41 | 0.33 | 0.73 | 0.67 |
| R1296 | **0.52** | 0.36 | **0.81** | 0.76 |
| R1271 | **0.78** | 0.66 | **0.79** | 0.77 |
| R1293 | **0.75** | 0.53 | **0.80** | 0.76 |
| R1261 | **0.75** | 0.56 | **0.77** | 0.72 |
| R1262 | **0.73** | 0.55 | **0.77** | 0.72 |
| R1211 | **0.74** | 0.50 | **0.81** | 0.75 |
| R1255 | **0.72** | 0.45 | 0.73 | 0.70 |
| R1256 | **0.57** | 0.39 | 0.61 | 0.56 |
| R1203 | **0.71** | 0.55 | **0.82** | 0.79 |
| **Avg.** | **0.65** | **0.49** | **0.77** | **0.73** |

#### S5 Target-wise performance of automated servers on CASP16

Table S5: Per-target **maximum** TM-score and lDDT for AlphaFold 3 (AF3-server, group 304) and trRosettaRNA2 (Yang-Server, group 052) on CASP16 targets (5 submissions per target). The last two columns report MSA depth (Deep MSA if Neff >130, otherwise Shallow) and CASP difficulty labels. For targets with mismatches between the provided FASTA sequence and the sequence extracted from the experimental/native structure, we compute scores only over the matched regions as determined by NW-align.

| **Target** | **AF3-server** | | **Yang-Server** | | **MSA** | **Difficulty** |
|---|---|---|---|---|---|---|
| | TM | lDDT | TM | lDDT | | |
| R1288 | 0.29 | 0.62 | 0.33 | 0.64 | Shallow | Hard |
| R1205 | 0.36 | 0.49 | 0.32 | 0.40 | Shallow | Hard |
| R1263 | 0.72 | 0.88 | 0.93 | 0.98 | Deep MSA | Easy |
| R1264 | 0.64 | 0.83 | 0.79 | 0.89 | Deep MSA | Easy |
| R1209 | 0.33 | 0.64 | 0.33 | 0.66 | Shallow | Hard |
| R1296 | 0.37 | 0.75 | 0.39 | 0.70 | Shallow | Hard |
| R1271 | 0.68 | 0.75 | 0.74 | 0.75 | Deep MSA | Medium |
| R1293 | 0.39 | 0.62 | 0.49 | 0.58 | Shallow | Medium |
| R1261 | 0.65 | 0.79 | 0.61 | 0.71 | Shallow | Medium |
| R1262 | 0.68 | 0.79 | 0.59 | 0.71 | Shallow | Medium |
| R1211 | 0.48 | 0.73 | 0.63 | 0.71 | Shallow | Medium |
| R1255 | 0.40 | 0.64 | 0.40 | 0.61 | Shallow | Medium |
| R1256 | 0.28 | 0.49 | 0.45 | 0.51 | Shallow | Medium |
| R1203 | 0.63 | 0.83 | 0.57 | 0.70 | Deep MSA | Medium |
| **Avg.** | **0.49** | **0.70** | **0.54** | **0.68** | | |

#### S6 Target-wise sampling performance of AlphaFold 3 and NuFold on CASP16

Table S6: Sampling performance without MSA. Columns 2–5 are averages over per-target maximum and means from 1000 sampled 3D structures using docker version of AlphaFold 3 with 200 random seeds (5 models generated per seed). Columns 6–9 show the corresponding metrics for samples generated from NuFold, using 200 random seeds (5 models per seed). For targets with mismatches between the provided FASTA sequence and the sequence extracted from the experimental/native structure, we compute scores only over the matched regions as determined by NW-align.

| **Target** | **AlphaFold 3** | | | | **NuFold** | | | |
|---|---|---|---|---|---|---|---|---|
| | TM-score | | lDDT | | TM-score | | lDDT | |
| | Max | Mean | Max | Mean | Max | Mean | Max | Mean |
| R1288 | 0.39 | 0.28 | 0.70 | 0.64 | 0.31 | 0.30 | 0.59 | 0.59 |
| R1205 | 0.40 | 0.29 | 0.49 | 0.45 | 0.41 | 0.39 | 0.44 | 0.44 |
| R1263 | 0.58 | 0.42 | 0.80 | 0.73 | 0.32 | 0.29 | 0.54 | 0.54 |
| R1264 | 0.55 | 0.42 | 0.79 | 0.73 | 0.31 | 0.28 | 0.54 | 0.54 |
| R1209 | 0.33 | 0.28 | 0.64 | 0.61 | 0.33 | 0.29 | 0.60 | 0.60 |
| R1296 | 0.45 | 0.31 | 0.73 | 0.69 | 0.34 | 0.28 | 0.65 | 0.65 |
| R1271 | 0.76 | 0.72 | 0.76 | 0.75 | 0.62 | 0.58 | 0.65 | 0.65 |
| R1293 | 0.40 | 0.31 | 0.63 | 0.60 | 0.33 | 0.31 | 0.59 | 0.59 |
| R1261 | 0.72 | 0.58 | 0.78 | 0.73 | 0.34 | 0.32 | 0.54 | 0.54 |
| R1262 | 0.71 | 0.58 | 0.78 | 0.73 | 0.35 | 0.33 | 0.55 | 0.55 |
| R1211 | 0.62 | 0.41 | 0.78 | 0.71 | 0.42 | 0.34 | 0.57 | 0.57 |
| R1255 | 0.49 | 0.33 | 0.66 | 0.61 | 0.34 | 0.27 | 0.52 | 0.52 |
| R1256 | 0.39 | 0.27 | 0.53 | 0.45 | 0.35 | 0.29 | 0.47 | 0.47 |
| R1203 | 0.66 | 0.55 | 0.84 | 0.82 | 0.48 | 0.45 | 0.65 | 0.65 |
| **Avg.** | **0.53** | **0.41** | **0.71** | **0.66** | **0.38** | **0.34** | **0.56** | **0.56** |

#### S7 Target-wise performance of RNAbpFlow on CASP16

Table S7: Per-target comparison of RNAbpFlow performance with and without the use of cross-distillation set during training. Average results on last row are based on 14 CASP16 RNA targets, presented in manuscript. For targets with mismatches between the provided FASTA sequence and the sequence extracted from the experimental/native structure, we compute scores only over the matched regions as determined by NW-align.

| | **RNAbpFlow (No distillation)** | | | | **RNAbpFlow (With distillation)** | | | |
|---|---|---|---|---|---|---|---|---|
| | **TM-score** | | **lDDT** | | **TM-score** | | **lDDT** | |
| **ID** | **Max** | **Mean** | **Max** | **Mean** | **Max** | **Mean** | **Max** | **Mean** |
| R1288 | 0.45 | 0.31 | 0.65 | 0.59 | 0.53 | 0.30 | 0.68 | 0.59 |
| R1205 | 0.36 | 0.24 | 0.47 | 0.34 | 0.40 | 0.28 | 0.50 | 0.36 |
| R1263 | 0.50 | 0.37 | 0.66 | 0.60 | 0.59 | 0.35 | 0.75 | 0.63 |
| R1264 | 0.49 | 0.36 | 0.69 | 0.61 | 0.60 | 0.33 | 0.77 | 0.63 |
| R1209 | 0.39 | 0.27 | 0.63 | 0.58 | 0.39 | 0.28 | 0.63 | 0.60 |
| R1296 | 0.47 | 0.34 | 0.75 | 0.67 | 0.50 | 0.32 | 0.77 | 0.70 |
| R1271 | 0.63 | 0.39 | 0.65 | 0.51 | 0.75 | 0.51 | 0.75 | 0.62 |
| R1293 | 0.55 | 0.41 | 0.66 | 0.56 | 0.56 | 0.41 | 0.67 | 0.61 |
| R1261 | 0.46 | 0.32 | 0.56 | 0.48 | 0.70 | 0.48 | 0.76 | 0.61 |
| R1262 | 0.46 | 0.32 | 0.57 | 0.49 | 0.68 | 0.47 | 0.77 | 0.61 |
| R1211 | 0.67 | 0.46 | 0.72 | 0.66 | 0.69 | 0.45 | 0.78 | 0.69 |
| R1255 | 0.57 | 0.36 | 0.60 | 0.56 | 0.64 | 0.39 | 0.66 | 0.61 |
| R1256 | 0.44 | 0.27 | 0.47 | 0.40 | 0.46 | 0.31 | 0.50 | 0.44 |
| R1203 | 0.63 | 0.44 | 0.72 | 0.66 | 0.71 | 0.47 | 0.83 | 0.75 |
| **Avg.** | **0.51** | **0.35** | **0.63** | **0.55** | **0.59** | **0.38** | **0.70** | **0.60** |

#### S8 Impact of base pair accuracy on CASP16

Table S8: Per-target score differences (Δ) (native − predicted base pairs) in TM-score and lDDT, alongside per-target p-values from paired Wilcoxon signed-rank tests on TM-score and lDDT distributions (1000 samples per target). Positive Δ values along with p-value $< 0.05$ indicate significant improvements when native base pair is used.

| | **TM-score** | | **lDDT** | | **Wilcoxon p-value** | |
|---|---|---|---|---|---|---|
| **ID** | **Max Δ** | **Mean Δ** | **Max Δ** | **Mean Δ** | **TM** | **lDDT** |
| R1288 | -0.10 | 0.05 | 0.10 | 0.15 | 2.5e−143 | 1.1e−165 |
| R1205 | 0.07 | 0.08 | 0.19 | 0.27 | 7.0e−152 | 2.6e−165 |
| R1263 | 0.18 | 0.27 | 0.12 | 0.19 | 4.8e−165 | 3.2e−165 |
| R1264 | 0.14 | 0.26 | 0.08 | 0.17 | 5.1e−165 | 2.1e−165 |
| R1209 | 0.02 | 0.05 | 0.10 | 0.07 | 4.0e−133 | 5.4e−166 |
| R1296 | 0.02 | 0.04 | 0.04 | 0.06 | 8.1e−83 | 5.6e−165 |
| R1271 | 0.03 | 0.15 | 0.04 | 0.15 | 6.0e−161 | 1.5e−165 |
| R1293 | 0.19 | 0.12 | 0.13 | 0.15 | 7.5e−162 | 1.6e−165 |
| R1261 | 0.05 | 0.08 | 0.01 | 0.11 | 3.5e−111 | 7.3e−161 |
| R1262 | 0.05 | 0.08 | 0.00 | 0.11 | 2.6e−105 | 7.2e−160 |
| R1211 | 0.05 | 0.05 | 0.03 | 0.06 | 1.3e−84 | 3.5e−163 |
| R1255 | 0.08 | 0.06 | 0.07 | 0.09 | 2.6e−102 | 5.3e−166 |
| R1256 | 0.11 | 0.08 | 0.11 | 0.12 | 1.6e−128 | 1.1e−165 |
| R1203 | 0.00 | 0.08 | -0.01 | 0.04 | 2.2e−66 | 6.1e−97 |

#### S9 Sampling statistics for native vs. predicted base pairs

Table S9: Comparison of sampling outcomes for native vs. predicted base pair inputs for 14 CASP16 targets.

| | **Native bp** | **Predicted bp** |
|---|---|---|
| % of at least one correct fold (TM-score > 0.45) | 85.71% | 71.43% |
| % of at least one correct fold (lDDT > 0.75) | 71.43% | 42.86% |
| % of correct folds (TM-score > 0.45) in ensemble | 56.48% | 24.66% |
| % of correct folds (lDDT > 0.75) in ensemble | 43.27% | 4.29% |

#### S10 Consistency between input and output base pairs

Table S10: Comparison of INF scores between input base pairs (provided during inference, either native or predicted) and output base pairs (those extracted from the corresponding predicted 3D structures using RNAView). Each row reports the average INF score, defined as INF = $\sqrt{\frac{TP}{TP+FP} \times \frac{TP}{TP+FN}}$, computed over 1000 predicted structures for each CASP16 target. Since multiple maps are provided as input during inference (3 maps for native and 15 maps for predicted, a consensus base pair list is constructed by including base pairs present in more than 50% of the input maps for the INF calculation in this experiment.

| **Target** | **Native bp (INF)** | **Predicted bp (INF)** |
|---|---|---|
| R1288 | 0.86 | 0.77 |
| R1205 | 0.83 | 0.68 |
| R1263 | 0.96 | 0.83 |
| R1264 | 0.96 | 0.83 |
| R1209 | 0.94 | 0.94 |
| R1296 | 0.96 | 0.94 |
| R1271 | 0.91 | 0.79 |
| R1293 | 0.87 | 0.82 |
| R1261 | 0.72 | 0.75 |
| R1262 | 0.72 | 0.75 |
| R1211 | 0.94 | 0.94 |
| R1255 | 0.95 | 0.87 |
| R1256 | 0.91 | 0.72 |
| R1203 | 0.95 | 0.89 |
| **Avg.** | **0.89** | **0.82** |

#### S11 Experimental base pair annotations differ across tools

Table S11: Total count of base pair annotations across RNAView (A), DSSR (B), and MC-Annotate (C) over 912 native RNA 3D structures collected from RNA3DB for CASP16 training set. Counts are summed over all RNAs. Percentages are relative to the union ($|A \cup B \cup C|$ = 37,877).

| Category | Count | % of Union |
|---|---|---|
| RNAView total ($|A|$) | 28,357 | 74.9 |
| DSSR total ($|B|$) | 21,045 | 55.6 |
| MC-Annotate total ($|C|$) | 35,030 | 92.5 |
| All three ($|A \cap B \cap C|$) | 19,622 | 51.8 |
| Union ($|A \cup B \cup C|$) | 37,877 | 100.0 |
